# Integrating opportunistic and structured non-invasive surveys with spatial capture-recapture models to map connectivity of the Pyrenean brown bear population

**DOI:** 10.1101/2022.05.13.491807

**Authors:** Maëlis Kervellec, Cyril Milleret, Cécile Vanpé, Pierre-Yves Quenette, Jérôme Sentilles, Santiago Palazón, Ivan Afonso Jordana, Ramón Jato, Miguel Mari Elósegui Irurtia, Olivier Gimenez

## Abstract

Connectivity, in the sense of the persistence of movements between habitat patches, is key to maintain endangered populations and has to be evaluated in management plans. In practice, connectivity is difficult to quantify especially for rare and elusive species. Here, we use spatial capture-recapture (SCR) models with an ecological detection distance to identify barriers to movement. We focused on the transnational critically endangered Pyrenean brown bear (*Ursus arctos*) population, which is distributed over Spain, France and Andorra and is divided into two main cores areas following translocations. We integrate structured monitoring from camera traps and hair snags with opportunistic data gathered after depredation events. While structured monitoring focuses on areas of regular bear presence, the integration of opportunistic data allows us to obtain information in a wider range of habitat, which is especially important for ecological inference. By estimating a resistance parameter from encounter data, we show that the road network impedes movements, leading to smaller home ranges with increasing road density. Although the quantitative effect of roads is context-dependent (i.e. varying according to landscape configuration), our model predicts that a brown bear with a home range located in an area with relatively high road density (8.29km/km^2^) has a home range size reduced by 1.4-fold for males and 1.6-fold for females compared to a brown bear with a home range located in an area with low road density (1.38km/km^2^). When assessing connectivity, spatial capture-recapture modeling offers an alternative to the use of experts’ opinion when telemetry data are not available.

## 1. Introduction

Habitat loss and fragmentation are a major concern for the conservation of animal populations (Fardila et al., 2017). Landscape structure can constraint the movement of individuals, their ability to find resources, to disperse, and to establish a home range (Fahrig, 2003). More generally, the presence of barriers can isolate populations and reduce their size, which can lead to the reduction of genetic diversity and ultimately affect population viability (Jackson and Fahrig, 2011). Landscape connectivity – “the degree to which the landscape facilitates or impedes movement among resource patches” (Taylor et al., 1993) – is increasingly included in conservation plans (Keeley et al., 2021).

Carnivores are usually the first species affected by the loss of connectivity because they live at low density and over large areas (Correa Ayram et al., 2016; Zeller et al., 2012). They are also flagship species, which increases stakeholder involvement in corridor projects (Beier et al., 2008). In particular, the transport infrastructure (e.g. road networks) has been a primary focus, because it causes direct mortality (i.e. roadkills), behavioral modifications (attraction or avoidance), or can act as physical barriers to movement (Coffin, 2007; Forman and Alexander, 1998).

A common approach to measure landscape connectivity is to build resistance maps. Resistance surfaces quantify the degree of potential flow through each cell given land cover types and using expert opinion or empirical data (Fletcher and Fortin, 2018). In practice, there is a trade-off between the information needed and the data available. Expert opinion is more subjective and less predictive of connectivity than biological data but it is often the only information available (Zeller et al., 2012). Telemetry and GPS data are the most informative data to study connectivity, because the process studied is movement (Zeller et al., 2018). However, these data are expensive to collect, can sometimes only be acquired at a coarse temporal resolution and often constitute a small proportion of the population monitored (Zeller et al., 2018, 2012).

Capture-recapture methods and, increasingly spatial capture-recapture (SCR) models, are the standard framework to monitor elusive populations from individual encounter data, like large carnivores (Royle et al., 2018). SCR models integrate a latent ecological process modeling the distribution of individuals and their activity centers, and a detection process accounting for heterogeneity in detectability by explicitly considering the distance between the traps and individual activity centers (Efford, 2004). The detection process of standard SCR models assumes circular home ranges, and therefore that the movement of individuals is homogeneous around the activity centers and not affected by landscape structure. When these assumptions are unlikely to be met, e.g. when habitat features restrict within home range movement, more complex SCR models can accommodate an ecological distance detection model. These models allow quantifying the impact of the landscape characteristics and estimating a resistance surface, as well as improving the estimation of space use patterns (Royle et al., 2013; Sutherland et al., 2015). Combined with the estimation of density, such SCR models allow an empirical quantification of connectivity based on spatial encounter histories from non-invasive detection data (Morin et al., 2017; Zeller et al., 2012). They model the distribution of activity center in space (i.e. second-order selection defined by Johnson, 1980) and estimate how landscape structure affects within home range movement and space use (i.e. third-order selection (Johnson, 1980)). Connectivity is measured at the scale of the home range in the sense of the “area traversed by the individual in its normal activities of food gathering, mating and caring for young” (Burt, 1943). Since SCR models can make use of monitoring data collected at the population level, and can account for imperfect detection, they can make population-level inferences (Royle et al., 2018).

In this study, we focused on the critically endangered Pyrenean brown bear (*Ursus arctos*) population, which is distributed over Spain, France and Andorra. Due to human persecution, the population was almost extinct in 1995 with only five individuals remaining in the western Pyrenees (Aspe and Ossau valleys) (Piédallu et al., 2019). Since 1996, eleven bears have been translocated from Slovenia in the western and the central Pyrenees to reinforce the population and avoid its extinction. The Pyrenean brown bear population is currently recovering with a minimum of 70 individuals detected in 2021 (Vanpé et al., 2022). The population is divided in two main core areas isolated with regard to exchange of females and located in areas with moderate human disturbance (Martin et al., 2012; Parres et al., 2020). Our objectives are to identify barriers that limits movement between resource patches and to evaluate how road fragmentation affects brown bear space use. The road network is assumed to impede brown bear movements, like it has been shown for other European populations (e.g. Slovakia: Skuban et al., 2017, Cantabrian: Mateo-Sánchez et al., 2014). However, the degree of avoidance to roads and the population-level consequences have never been quantified. We used transnational non-invasive genetic sampling data across France and Spain to draw population-level inferences about connectivity. To do so, we accounted for imperfect and heterogeneous detections by building a SCR model with non-invasive genetic sampling data obtained from the structured and opportunistic monitoring of the population.

## 2. Material and Methods

### 2.1. Study area

The transnational study area (36,600km^2^) is located at the border between Southwestern France (6 counties), Northeastern Spain (3 autonomous regions) and Andorra, and encompasses the entire range of the Pyrenean brown bear population (Figure 1). Elevation ranges from 0 to 3,400m and is characterized by large valleys and steep mountains. The study area is composed at 41% of forests with deciduous (dominant beech *Fagus sylvatica*) and coniferous trees (dominant silver fir *Abies alba*) (Martin et al., 2012). Between forest patches there are large open areas with shrubs such as rhododendron (*Rhododendron ferrugineum*) and wild blueberry (*Vaccinium myrtillus*) above 1,800m. The landscape is also shaped by human activities. Traditional pastoral activity, mostly sheep, occupies pastures at higher altitudes from June to October. Other human activities are forestry and recreational activities (e.g., hiking, hunting, skiing). Human population (mean = 67 inhabitants/km^2^), and the road network (mean = 3.8 km/km^2^) are mostly concentrated in the valleys. The Pyrenean Mountains are framed by highways, and few primary roads cross perpendicularly the massif to link France and Spain (Figure A2).

**Figure 1.**
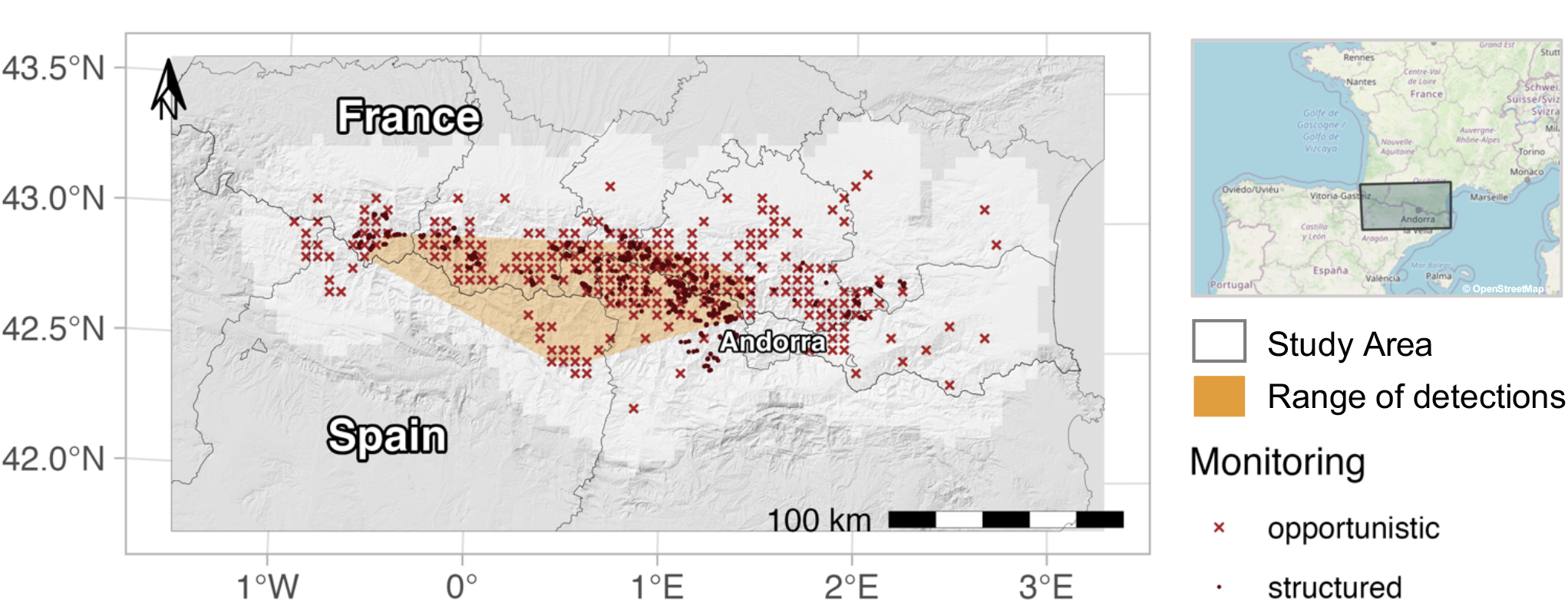
Study area, minimum convex polygon to represent range of detections and locations of traps for the monitoring of the brown bear population in the Pyrenees in 2019.

### 2.2. Data collection

#### 2.2.1. Structured monitoring

Three surveys were conducted in 2017, 2018 and 2019 from May to November. We assumed that the population was demographically closed since births take place in winter and the Pyrenean population is isolated from other populations. We used a structured sampling design, and we set up two types of traps, DNA hair snags alone or combined with camera traps. These traps were distributed in the areas known to be regularly occupied by bears in France and in Catalonia. Camera traps were essentially set up to detect reproductions, while hair snags allowed the genetic identification of individuals. Some individuals with distinctive marks (GPS collar, ear mark, distinctive spot) could also be identified on camera traps. Hair snags were baited with beech tar. We visited the stations at least once a month, and twice for some traps in France, in May, June and September. More details about the monitoring protocols can be found in Vanpé et al. (2022).

#### 2.2.2. Opportunistic monitoring

We also used opportunistic detection data consisting in all the genetic samples collected following a depredation event on livestock or beehives. We did not consider other opportunistic signs of presence, because we were not able to define a corresponding search effort. In order to match the study period defined for the structured monitoring, we only considered those collected from May to November. The trap array can be assimilated to the distribution of livestock in the Pyrenees. It is recommended that aggregated traps be spaced 1.5 σ apart (σ is the scale parameter estimated in spatial capture-recapture model), or approximately 5km for bears in the Pyrenees (Milleret et al., 2018). To match opportunistic detection data to a flock, we defined a 5×5 km grid and consider a cell to be active if at least one depredation event occurred between 2010 and 2020. Then, every sample collected was attributed to the center of the cell, making all centers of active cells opportunistic traps.

#### 2.2.3. Brown bears individual identification

The individual identification of bears by genetic analysis of the samples (hair) consisted in the amplification and identification of genetic markers. We obtained species and genetic lineage by mitochondrial DNA analysis. We determined individual identity using a multiple tubes Polymerase Chain Reaction (PCR) approach with 13 microsatellite markers (Vanpé et al., 2022). Sex was identified by combining three markers present on the sex chromosomes (Vanpé et al., 2022). When a sample could not be attributed to an individual, it was discarded from the analyses. In total, 156 samples were genetically identified in 2017, 110 in 2018, and 200 in 2019, which corresponded to 31, 31, and 33 different individuals, respectively (Table A1).

### 2.3. Statistical modelling: Spatial capture-recapture model

We estimated bear density and landscape connectivity simultaneously with spatial capture-recapture (SCR) models. These models combine 1) a detection process to account for the imperfect detection of individuals from individual encounter history and 2) a spatial point process to model the distribution of individuals in space through the estimation of their activity center’s location (Efford, 2004; Royle et al., 2013). Here, we used a multi-session sex-structured SCR model (Sutherland et al., 2019). This model provides a framework to account for variation in densities between males and females and between different monitoring sessions. First, we fitted the SCR model to the structured survey data and accounted for heterogeneity in detection. Second, we integrated the opportunistic data, and evaluated the added value of data integration to improve inference about spatial heterogeneity in density, detection and resistance. Finally, we quantified the impact of roads on the shape of brown bear home range, using non-Euclidean distance SCR models (Sutherland et al., 2015). Parameters were estimated by maximum likelihood implemented in R using the package oSCR (Sutherland et al., 2019).

#### 2.3.1. Baseline detection probability

SCR models are based on encounter history data *y_ijk_* of an individual *i* at trap *j* during capture occasion *k*:

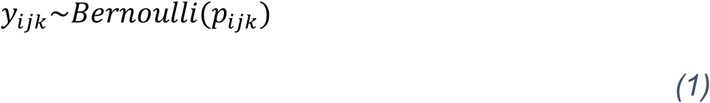

where the detection probability, *p_ijk_* describes the probability to detect an individual *i* at trap *j* during sampling occasion *k*. We assumed that this probability decreases with the distance between the activity center and the trap. We modeled this probability with a half-normal detection function:

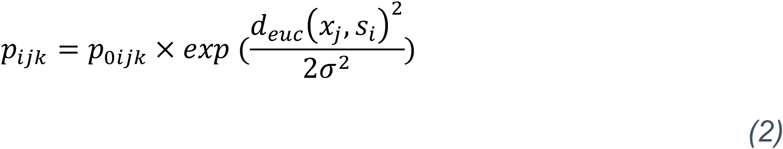

where *d_euc_(x_j_,s_i_)* is the Euclidean distance between trap *x_j_* and activity center *s_i_*. The baseline detection probability *p_0ijk_* is the detection probability when an individual’s activity center is exactly on the trap (i.e. *d_euc_(x_j_,s_i_) = 0)*.

We considered six variables to explain variation in *p_0ijk_* (Table 1):

**Table 1.** Description of the SCR model parameters and variables considered

| Parameter | Notation | Meaning |
| --- | --- | --- |
| $p_0$ | $b$ | Behavioral response to traps |
| | $m + m^2$ | Quadratic effect of the sampling occasion |
|  | session | Year (2017, 2018, 2019) |
|  | sex | Sex (female, male) |
|  | effort | Search effort: country and number of visits in the station per month (1 visit in France, 2 visits in France, 1 visit in Spain) |
|  | trap | Type of traps: structured (hair snag alone or hair snag combined with camera trap), or opportunistic (center of 5 km grid surrounding depredations since 2010) |
| sigma | sex | Sex |
| density | elevation | Mean elevation over 200 km<sup>2</sup> |
|  | ruggedness | Mean terrain ruggedness index over 200 km<sup>2</sup> |
|  | human density | Log mean human density over 200 km<sup>2</sup> |
|  | forest cover | Mean forest cover over 200 km<sup>2</sup> |
| movement | road density | Length of all types of roads (in km) by cells of 6.25 km<sup>2</sup> |

1. The behavioral response (**b**) is a binary individual covariate that differentiates the first detection of an individual at a given trap (b=0) from the following ones. This persistent response accounts for trap happiness, because hair snags are baited and individuals detected once seems more likely to be detected afterwards.
2. The sampling occasion (**m**) is defined as a month from May to November (m = 1,.., 7). To acknowledge that bears are more likely to be detected in the summer, we considered a quadratic effect of the sampling occasion (**m + m^2^**).
3. The search effort (**effort**) is a three-level factor. It is defined according to the country where the trap is placed (France or Spain) and the number of visits performed per month to collect genetic material. Our hypothesis is that the greater the number of visits at a station, the greater are the chance to obtain DNA material on the trap of sufficient quality to allow identification of the individual (De Barba et al., 2010). The search effort at opportunistic traps was assumed to be equivalent to traps in the structured monitoring that were visited once per month, and depended of the country where the depredation occurs.
4. The type of trap (**trap**) describes whether the trap consists of a hair snag alone, a hair snag combined with a camera trap, or an opportunistic trap. Hair snag combined with a camera trap is supposed to have higher detection probability, because hair founded in front of a camera trap are identified in priority.
5. The session (**session**), here define by the year (2017, 2018, 2019), accounts for annual variations in detection probability.
6. The sex covariate (**sex**) is considered to model behavioral differences (e.g., females with cubs avoid areas where males are present, and males disperse over large distances) (Swenson et al., 2000).

#### 2.3.2. Scale parameter

The scale parameter σ controls the shape of the detection function. The larger σ is, the slower the detection probability decreases with distance from the activity center. We assumed that σ varied with sex because males are known to have larger home range than females (Swenson et al., 2000). In preliminary analyses, we noticed that two males, Néré and Goiat, were detected over half of the study area. As we could not assume that they had the same home range size as the other males and to avoid bias in σ estimate, we dropped these individuals from our analyses. From the estimation of σ in the case of a half-normal detection function, it was possible to estimate the size of the home range according to the relation: 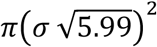 (Royle et al., 2014).

#### 2.3.3. Spatial variation in density

Activity centers *s_i_* can be uniformly and independently distributed in the spatial domain *S* (set of potential positions of the activity centers of detectable individuals) with constant intensity, or the intensity can vary according to landscape variables. This spatial domain is defined from the traps around which we add a buffer zone (i.e. a zone large enough to encompass all activity centers of individuals which can be detected by the trap array) (Sutherland et al., 2019). We defined a buffer of 25km around the stations (approx. 3 and 4σ) and used a 5km resolution of the spatial domain.

We selected four habitat variables (Figure A1) to model spatial variation in density of bears in the Pyrenees:

1. Forest cover is obtained from the Corine Land Cover (European Environment Agency, 2018) forest data (deciduous, coniferous, and mixed forests) for France and Spain at 100m and the Andorra habitat map of 2012 (Institut d’Estudis Andorrans, 2012) at 250m resolution.
2. Elevation is defined at 90m resolution (Shuttle Radar Topographic Mission, 2018). High elevation generally correlates with areas of low human presence and low food resource availability.
3. Ruggedness is the average of the absolute values of elevation differences between the focal cell and the eight surrounding cells.
4. Human density (Columbia University, 2018) is intended to inform the model about human activities that bears seek to avoid (Swenson et al., 2000).

To ensure that each variable described the habitat at the scale of the bear home range and not just at a given point, we averaged each habitat variable with a sliding window of 200km^2^. The size of the window corresponds to the average home range size of an adult female bear estimated in preliminary analyses. We scaled the three first variables and log-transformed and scaled human density to obtain four rasters of habitat at 5×5 km^2^ resolution.

### 2.4. Estimation of road resistance on connectivity from detection data

The shape of individuals’ home ranges depends on the distribution of resources in the landscape as well as constraints on their movements, like the presence of roads (Dahle and Swenson, 2003, Proctor et al. 2019). Home ranges can be irregular, asymmetric, and non-stationary (i.e. varying with location) (Royle et al., 2013). We accounted for the impact of landscape structure on movement using the ecological distance. This distance, involved in the calculation of p_ijk_ (Eq. 2), is based on the least cost path distance instead of the Euclidean distance. Given this discrete landscape v, the distance between two points can be represented by a sequence of m steps connecting cells denoted v_1_, v_2_, …, v_p_. We computed the cost of joining two points through this path, and through all possible paths 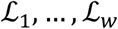. The least cost path is defined as the minimum of these paths:

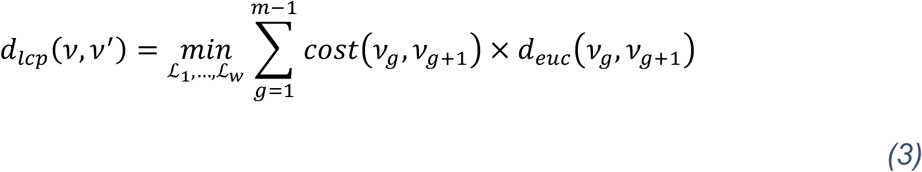

where 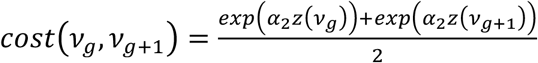

The cost of using a cell is estimated by the resistance parameter α_2_, which is estimated with the maximum likelihood method and by using the value taken by the variable of interest *z(v_p_)* in the cell *v_p_*. The estimated resistance parameter reflects the extent to which a given habitat variable facilitates (α_2_ < 0) or impedes (α_2_ > 0) the movement of individuals across the landscape (Royle et al., 2013; Sutherland et al., 2015). When α_2_ = 0 (i.e. the resistance is null) we have *cost(0) = 1* and the distance is exactly Euclidean.

The probability that an individual with its activity center *s_i_* uses a cell *su* in *S*, noted *Pr(g[s_u_,s_i_])*, can be calculated from the estimated parameters and *p_0_ = 1*. We distinguished cell use (g) from imperfect detection (y) (Sutherland et al. 2015). Given an activity center location, we computed the 95% kernel of the utilization distribution to estimate the home range size in a particular landscape context. The potential connectivity, denoted PC, represents the total utilization assuming that one individual activity center is located in each cell of the habitat. In other word, PC illustrates the number of individuals reaching a cell given that individuals are evenly distributed in the landscape. In the oSCR package, the matrix recording the least-cost path between all cells v, is computed by the Dijkstra algorithm implemented in the *gdistance* package (van Etten, 2012). Realized densities and potential connectivity can be combined to obtain a realized map of connectivity called density weighted connectivity (DWC) for males and females since their home range size are estimated separately (Morin et al., 2017).

To determine the impact of roads on connectivity, we built a landscape variable considering the length of all roads defined in OpenStreetMap from motorway to track at a resolution of 2.5km (Figure A2). This covariate is not correlated (Pearson r < 0.7) to the four covariates considered as related to density (Table A2).

### 2.5. Model selection

To avoid having to test too many models we conducted a hierarchical selection. The best model was selected according to the Akaike information criterion (AIC; Burnham and Anderson, 2002).

#### 2.5.1. Model heterogeneity in detection probability

Firstly, we focused on the detection probability and we tested all combinations of behavioral, sampling occasion, session and sex on p0 only with data from the structured monitoring. We included type of trap and search effort as additive linear effects on p0 and a sex effect on σ in each model. At the same time, we also tested whether density was constant between 2017 and 2019 or varied between sessions. In this step, we compared 32 models.

#### 2.5.2. Model spatial variation in density with habitat and integration of opportunistic data

Secondly, we selected the variables explaining best spatial variation in density according to the AIC. In case several models had a better AIC than the model with uniform density, we also tested their additive effect if they were not correlated (Pearson r < 0.7). We conducted the same model selection procedure on models considering only structured detection data and on models with both structured and opportunistic detection data, resulting in 10 additional models to compare. We hypothesized that the integration of opportunistic data could improve inferences of spatial variation in density (Tenan et al. 2017). In addition to providing more detections, it also improved the spatial coverage of the survey as opportunistic traps area also located in areas with low density or absence of bears, contrary to the structured survey that was located only in areas regularly occupied by bears (Figure 1).

The model best supported by the data then served as the basis for quantifying connectivity. Finally, we used AIC to evaluate whether the Euclidean model performed better than the non-Euclidean one (Sutherland et al., 2015).

## 3. Results

### 3.1. Heterogeneity in detection probability in the structured monitoring data

The eight best models (AIC differences with the best model: ΔAIC < 6) modelling heterogeneity in detection probability included a behavioral response and a quadratic effect of time. Because estimates were close, we used in the next step the model with the lowest AIC (Table A3). In this model, density was constant in the three monitoring sessions. The baseline detection probability, p0, was maximum in June and July, close to zero in October and November and was higher if the bear had already been detected once. The baseline detection probability was also higher if the trap was composed of a camera trap and hair snags and was visited twice per month in France (Figure A3). The estimated scale parameter (*σ*) was larger for males, 7.31km (CI_95_ = [6.56, 8.13]), than for females, 4.89km (CI_95_ = [4.35, 5.49]), as we anticipated (Figure A4). According to the relationship 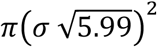 and the assumption of circular home range we found that the estimated 95% home range size was 252.1km^2^ (CI_95_ = [179.0, 355.1]) for females, and 1433.3km^2^ (CI_95_ = [1056.5, 1944.5]) for males.

### 3.2. Spatial variation in density

When considering only detection data from the structured monitoring, none of the models that accommodated spatial covariates on density performed better than the null model (Table A4. A). In contrast, after integrating the opportunistic detection data, the model with an additive effect of ruggedness and human density was considered as the best model (Table A4. B). Bear density was negatively correlated with human density (Figure 2D), and positively correlated with ruggedness (Figure 2C). When there was on average one inhabitant/km^2^ over the 200km^2^ surrounding, brown bears density was estimated maximum at 0.015 (CI_95_ = [0.007,0.029]) and 0.009 (CI_95_ = [0.005,0.018]) respectively for females and males (Figure A5).

**Figure 2.**
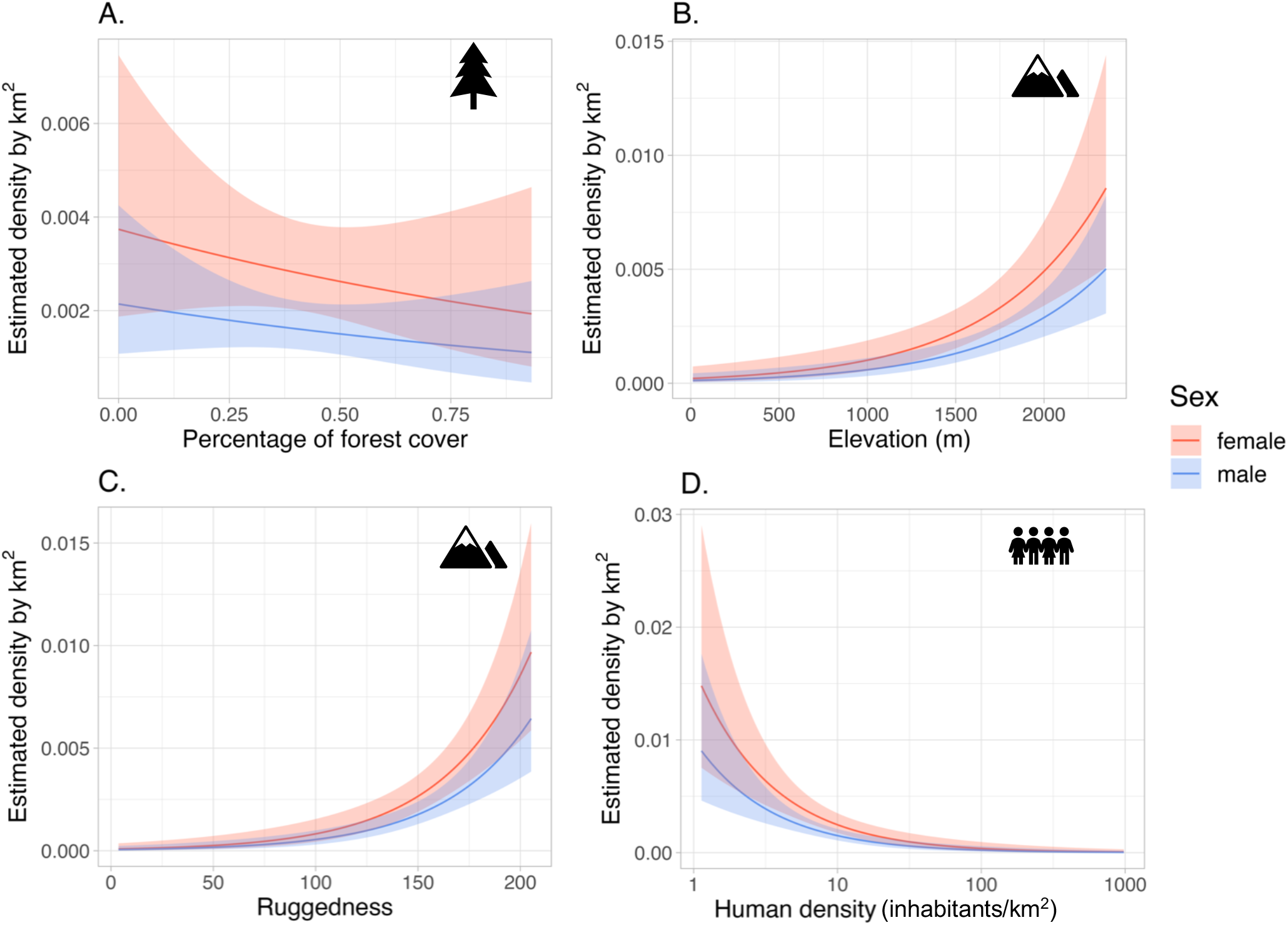
Estimated density of male (blue line) and female (red line) Pyrenean brown bears per km^2^ as a function of four habitat covariates: (A) percentage of forest cover, (B) elevation (m), (C) ruggedness and (D) human density (mean number of inhabitants per km^2^ over 200 km^2^). The curves represent the estimated values and the shaded zone the confidence interval at 95%.

### 3.3. The influence of roads on connectivity

The use of non-Euclidean distance SCR model, with the density of roads as a spatial covariate, strongly improved the AIC of the model (AIC = 3995.5). The estimated resistance parameter *α*_2_ = 0.428 (CI_95_ = [0.12,0.74]) was positive (Table A5), meaning that the density of roads impeded movement of brown bear. We selected two representative points of high and low road density within the brown bear range. In an area with high density of roads (i.e. 8.29km/km^2^, near Vielha in Val d’Aran, Spain), the model predicted 95% home range size to be 736.5km^2^ for males and 321.1km^2^ for females. Conversely, in an area with low density of roads (i.e. 1.38km/km^2^, near Couflens in Ariège, France), the model predicted 95% home range size to be 1052.7km^2^ for males and 549.9km^2^ for females (Details about the variation of home range size according to road density are available in the supplementary material (Figure A6 & A7)).

## 4. Discussion

Using a non-Euclidean SCR model (Morin et al., 2017; Sutherland et al., 2015) and multiple sources of non-invasive sampling data, our study provided evidence that roads acted as a barrier to the movement of brown bears in the critically endangered Pyrenean population. The home range size of brown bears is smaller with increasing density of roads, which can impede the connectivity and ultimately limit access to resources patches. In addition, we showed the importance of collecting and integrating opportunistic sampling data in SCR models to reveal ecological patterns, such as spatial variation in density and connectivity.

### 4.1. Spatial variation in density

Bear density was not uniformly distributed over the study area and decreased with human density and increased with ruggedness. Rugged terrain can be considered as refuge areas as they are generally remote and characterized by terrain with high elevation. These results are consistent with previous studies of this population (Martin et al., 2012; Piédallu et al., 2019), and also on other bear populations (e.g. Cantabrian: Mateo-Sánchez et al., 2014; Andean: Morrell et al., 2021). These selected variables are habitat descriptors that correlate relatively well with the characteristics of the two cores areas where the bears from Slovenia were translocated and established their home range, but are not necessarily synonymous of high-quality habitat (Parres et al., 2020). These results cannot therefore be used to infer the future distribution of the population. This also means that other uncolonized habitat which is not characterized by high ruggedness and low human density may support the presence of bears.

### 4.2. Integration of opportunistic monitoring

In the Pyrenees, the structured monitoring is restricted to the area of regular bear presence, while the opportunistic sampling can, in theory, occur anywhere where domestic animals are located and where bear depredation can occur. The integration of opportunistic detection data using a detector grid located almost continuously through the population range (Figure 1) allowed us to have traps with no detection of any individual in areas with low or sporadic bear presence. Obtaining information in such areas was very important to detect spatial variation in density and likely the reason why we could only detect association between density and habitat variables after integrating opportunistic samples (Sun et al., 2019). By integrating multiple data sources into SCR models, we provided density maps of the Pyrenean brown bear population across its entire distribution range, which was not possible by considering the structured monitoring only.

Brown bear is an elusive species living at low density over large areas (Swenson et al., 2000). SCR models account for imperfect and heterogeneous detection probability, which varies across time, space and individual characteristics (Efford, 2004; Sutherland et al., 2019). The baseline detection probability varied during the year according to the ecological characteristics of the species, with higher detectability in June during the mating season, and in the summer when their frequency of depredation events usually picks, before it decreased to be close to zero in October and November during hyperphagia when bears can stay on the same feeding area for several days before hibernation (Swenson et al., 2000). The baseline detection probability was also higher when a bear was at least detected once. This behavioral effect accounted for the trap happiness, because traps are baited, and also because some individuals seem to have a higher detection rate than others. Considering heterogeneity in detectability was important in order to obtain unbiased abundance estimates.

Usually, the monitoring of large carnivores is composed of different types of survey to maximize the number of individuals detected (e.g. Sollmann et al., 2013). Here we considered two types of surveys: the structured monitoring (i.e. hair snags and camera traps) and the opportunistic monitoring (i.e. biological samples gathered after a depredation). Camera traps alone do not permit the identification of bears. They inform on the presence of cubs, the date to which a bear left hair on the traps, and whether several bears use the same hair snag between two visits. Usually, opportunistic detection data are not considered even though they allow the capture of individuals missed or the spatial recapture of bears already captured by structured monitoring, and can improve the spatiotemporal extent of inferences (Sun et al., 2019).

The main obstacle to the use of opportunistic detection data is the difficulty to obtain robust data on the spatial variation in search effort because the detection process is opportunistic and therefore not recorded (Zipkin et al., 2021). Consequently, we had to consider a spatially homogeneous opportunistic search effort. In the spirit of Tenan et al. (2017), we checked for data consistency by comparing parameter estimates in the model integrating or not opportunistic data (Figures A3 & A4). In our case, the baseline detection probability at structured monitoring traps (hair snags alone and hair snags combined with a camera trap), as well as the scale parameter, did not change after the integration of opportunistic detection data (Figure A3 & A4). In the integrated model, the baseline detection probability increased with the number of visits (i.e., the search effort). When visits are too sparse, the DNA may be degraded, in too small quantity, or the sample may be composed of DNA originated from several bears (De Barba et al., 2010), leading to genotyping issues. Moreover, some genetic material could not be analyzed for financial reasons. Failure to identify the bear associated with a detection may lead to overestimation of densities and underestimation of p_0_ (Royle et al., 2014). Here the population is well known and individuals are recaptured frequently. This process is unlikely to happen because we estimated a population size close to actual population counts (Vanpé et al., 2022).

### 4.3. The influence of roads on connectivity

We found that road density impeded the movement of brown bear. In an area with low road density (1.38km/km^2^), home range is 1.4 times larger for males and 1.6 times larger for females than in an area with high road density (8.29km/km^2^). This shows that landscape fragmentation and roads can act as barrier and as a limiting factor for the distribution, and space use of individuals (Coffin, 2007). This situation is similar to the case of the Cantabrian brown bear population which were divided into two cores areas and suffered from low genetic diversity (Pérez et al., 2009). Spanish managers focused on restoring connectivity by creating a corridor between the two nuclei (tree plantations, crossings) and these actions now show encouraging results (i.e. improve demographic and genetic exchanges) (Gonzalez et al., 2016). The area near Andorra, the Mediterranean coast between Narbonne and Gerona and the north-west part of the study area near Pau and Tarbes have a low potential connectivity, suggesting that connectivity might be challenging to achieve in this part of the Pyrenees. Roads from Saint-Gaudens to Vielha limit connectivity in the west of the centro-oriental core, which may limit connectivity between the two cores (Figure 3). Note that we consider that all roads have the same weight even though forestry roads do not have the same ecological effect as highways, that is roads with a high traffic volume or which are fenced are less crossed (Skuban et al., 2017). During preliminary analyses relying only on French data different characterizations of the road network were explored (distance to main roads (i.e. motorway, trunk and primary roads) at 2.5km, density of paved roads (i.e. motorway, trunk, primary, secondary and tertiary roads) at 1km, density of all roads at 1km). The model considering the density of all roads had the lowest AIC and was retained for further analyses (Table A6). However, this choice of implementation gives more importance to urban areas, which have higher road density. Moreover, our model remains correlative and other variables correlated with road density, might be ecologically responsible of the resistance to movement (Royle et al., 2013). In the Pyrenees, roads are concentrated in valleys, where there are villages, train track, semi-canalized rivers, human activities, higher traffic volume and lower food availability (Martin et al., 2012). These factors can also shape brown bear movements (Proctor et al., 2019, Skuban et al., 2017, Swenson et al., 2000).

**Figure 3.**
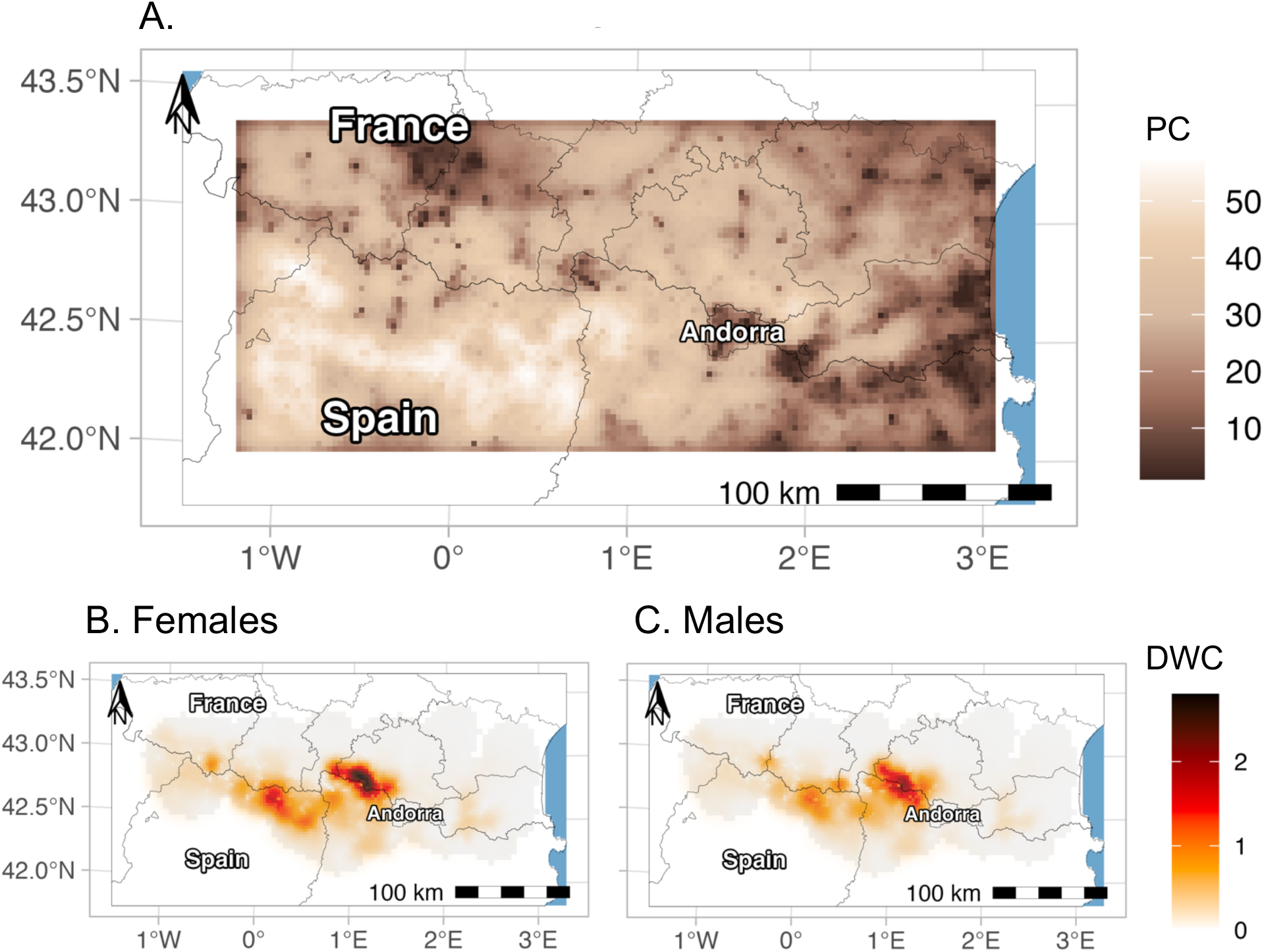
Maps of metrics of connectivity for the Pyrenean brown bear population in 2019. (A) Potential connectivity (PC) for both sex and density weighted connectivity (DWC) for (B) females only and (C) males only. The grey shaded zone in (B) and (C) represents the study area.

Density weighted connectivity (DWC) provides a realized measure of connectivity at the population scale (Morin et al., 2017). DWC map showed two cores (Figure 3). The larger one was in the centro-oriental part of the Pyrenees at the boundary between Ariège and Catalonia and the second is in the western Pyrenees (Piédallu et al., 2019). However, we also estimated a high DWC in the Ordesa y Monte Perdido National Park, because the model predicted the presence of undetected individuals. The model likely predicted the presence of bears since the habitat characteristics in this area was associated with high bear density (very low human density (between 0 and 3 inhabitants/km^2^) and steep slopes (Figure 3)), but with a low recorded search effort. The spatial capture-recapture framework including an ecological distance allows the estimation of a resistance surface based on encounter history data (Sutherland et al., 2015). Compared to occupancy models with unknown individual identity (Howell et al., 2018, Vasudev et al., 2021), SCR models that use known individual identity, encounter data offer the advantage to mechanistically model the influence of covariates on individual movement. This framework offers an efficient and cost-effective alternative to the use of expert opinion or telemetry data to quantify connectivity (Royle et al., 2018; Zeller et al., 2012). However, if GPS or telemetry data are available, they can be integrated with capture-recapture data to provide more precise estimate of movement and inform the estimation of the scale parameter (Dupont et al., 2021).

Euclidean SCR models assume a half-normal detection function, meaning that the estimated home range is circular, regular and symmetric, which is biologically unrealistic and lead to biased estimates of density (Royle et al., 2013). Also, the consideration of landscape characteristics explaining variation in the size and the shape of home ranges provided us with a more realistic measure of space use. Our estimates of home range size are comparable to what was found for other European populations (e.g. Southern Scandinavia: males [314, 8264] km^2^, and females without cubs [81, 999] km^2^ (Dahle and Swenson, 2003) or in Trento province in Italia [34,1813] km^2^ (Preatoni et al., 2005)). However, we supposed that all males and all females had a similar home range size, which can be a source of bias if some individuals display higher movement abilities. Here, we discarded such individuals from the analysis, even though their contribution to connectivity could be important. Although the ecological reason for this outlier behavior remains unclear, finite mixture models associated with spatial capture-recapture framework could be used to capture this heterogeneity and the contribution of these individuals to connectivity be quantified.

The SCR framework does not explicitly integrate movement like other connectivity models based on GPS and telemetry data, which precludes from modeling trajectories between captures (Zeller et al., 2018). The SCR framework models the habitat used by an individual during a year around its activity center, therefore integrating concepts (e.g. resistance surface) consistent with the landscape connectivity theory (Royle et al., 2013). Moreover, the density weighted connectivity provides a functional metric of connectivity which is not simply the inverse of the habitat suitability (Keeley et al., 2016; Morin et al., 2017).

### 4.4. Conservation implication

Spatially explicit estimator of connectivity and density are key to the conservation of recovering populations. However, the biological movement data needed to obtain such estimator are often lacking or difficult to collect in sufficient quantity (Zeller et al., 2012). SCR models with ecological distance provide a framework to integrate structured and opportunistic detection data often collected in monitoring programs on carnivores. SCR models also permit avoiding double counting of individuals that live on both sides of a boundary, as it is often the case in transnational populations (Bischof et al., 2016), and therefore population size overestimation. Overall, SCR models enable ecologists without GPS data to identify problematic areas that limit movement and to estimate a resistance surface from encounter data. These maps based on the probability of space usage and population density provide spatial information for managers to place wildlife crossing for example (Morin et al., 2017; Royle et al., 2013) and corridors, although we acknowledge that the implementation of corridors should consider other species requirements (e.g. dispersal abilities, or migratory behavior) (Beier et al., 2008).

## Appendices

**Table A1.** Summary of search effort for the three years monitored in the Pyrenees Mountains (France, Spain, Andorra) and the two monitoring methods (structured, opportunistic): the number of traps (hair snags, hair snags and camera traps or center of 5 km grid surrounding depredations since 2010), number of independent detections (i.e. number of independent bear detections per trap and per sampling occasion defined as a month), number of unique individuals detected (i.e. number of unique bears identified per survey), average detections (average numbers of time a bear has been independently detected during a survey), and average spatial detections (average numbers of stations a bear has been independently detected during a survey)

| Year | Monitoring | No. of traps | No. of independent detections | No. of unique individuals | Average detections | Average spatial detections |
| --- | --- | --- | --- | --- | --- | --- |
| 2017 | Structured | 654 | 71 | 23 | 3.09 | 2.65 |
| 2017 | Opportunistic | 246 | 29 | 17 | 1.71 | 1.59 |
| 2018 | Structured | 696 | 46 | 21 | 2.19 | 1.95 |
| 2018 | Opportunistic | 246 | 43 | 21 | 2.05 | 1.81 |
| 2019 | Structured | 692 | 95 | 26 | 3.65 | 3.08 |
| 2019 | Opportunistic | 246 | 52 | 19 | 2.74 | 2.21 |

**Table A2.**
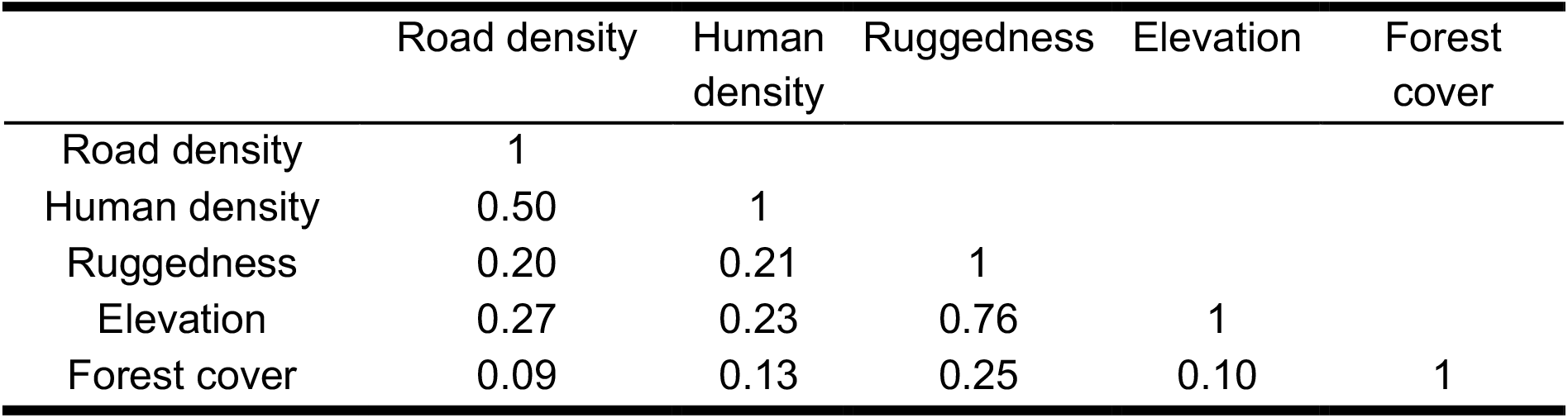
Matrix of Pearson correlation between forest cover, elevation, ruggedness, road density and human density aggregated at a resolution of 2.5km.

|  | Road density | Human density | Ruggedness | Elevation | Forest cover |
| --- | --- | --- | --- | --- | --- |
| Road density | 1 |  |  |  |  |
| Human density | 0.50 | 1 |  |  |  |
| Ruggedness | 0.20 | 0.21 | 1 |  |  |
| Elevation | 0.27 | 0.23 | 0.76 | 1 |  |
| Forest cover | 0.09 | 0.13 | 0.25 | 0.10 | 1 |

**Table A3.** Eight best models and the null model among the 32 tested, ranking according to their AIC to account for heterogeneity in the detection process considering only data from structured monitoring. D is density, p_0_ is the baseline detection probability, K is the number of estimated parameters, AIC is the Akaike Information Criterion and ΔAIC is the difference between the AIC of each model and the model with the lowest AIC. Each model considers the type of trap (hair snag alone or combined with a camera trap) and the search effort (number of visits at a trap per month and country) on the baseline detection probability p_0_ and a sex effect on the scale parameter σ. **Notation**: *b* is the behavioral response to traps (binary individual covariate that differentiates the first detection of an individual (b=0) from the following ones (b=1)), *m+m2* corresponds to the quadradic effect of the month, *session* is the year when the study was conducted and *sex* is the sex of the individual (female or male)

| D | $p_0$ | K | AIC | $\Delta AIC$ |
| --- | --- | --- | --- | --- |
| 1 | $b + m + m^2$ | 11 | 2528.90 | 0.00 |
| 1 | $b + m + m^2 + \text{session}$ | 13 | 2528.94 | 0.04 |
| 1 | $\text{sex} + b + m + m^2 + \text{session}$ | 14 | 2530.71 | 1.81 |
| 1 | $\text{sex} + b + m + m^2$ | 12 | 2530.86 | 1.96 |
| session | $b + m + m^2$ | 13 | 2532.58 | 3.68 |
| session | $b + m + m^2 + \text{session}$ | 15 | 2532.67 | 3.77 |
| session | $\text{sex} + b + m + m^2 + \text{session}$ | 16 | 2534.40 | 5.50 |
| session | $\text{sex} + b + m + m^2$ | 14 | 2534.54 | 5.64 |
| 1 | 1 | 8 | 2607.44 | 78.54 |

**Table A4.** Candidate models ranking according to their AIC to account for variations of bear density according to four habitat covariates. (A) considers only structured data and (B) considers all data (structured and opportunistic). D is the density, K is the number of estimated parameters, AIC is the Akaike Information Criterion and ΔAIC is the difference between the AIC of each model and the model with the lowest AIC. Each model considers that the baseline detection probability p_0_ is a linear function of the behavioral response to traps (b), the quadratic effect of the month (m + m^2^), the type of trap (trap) (i.e. hair snag alone, hair snag combined with a camera trap, or the center of the active cell in the opportunistic grid at 5 km resolution), and the search effort (effort) (i.e. depends of the country where the trap is located and the number of visits per month). The scale parameter σ is sex dependent. (p_0_ ~ b + m + m^2^ + trap + effort) (σ ~ sex)

| D | K | AIC | $\Delta AIC$ |
| --- | --- | --- | --- |
| 1 | 11 | 2528.90 | 0.00 |
| percentage of forest cover | 12 | 2530.40 | 1.50 |
| ruggedness | 12 | 2530.58 | 1.68 |
| human density | 12 | 2530.67 | 1.77 |
| elevation | 12 | 2530.86 | 1.96 |

B.
| D | K | AIC | $\Delta AIC$ |
| --- | --- | --- | --- |
| ruggedness + human density | 14 | 4009.40 | 0.00 |
| ruggedness | 13 | 4018.51 | 9.11 |
| human density | 13 | 4028.38 | 18.98 |
| elevation | 13 | 4031.27 | 21.87 |
| 1 | 12 | 4055.00 | 45.60 |
| percentage of forest cover | 13 | 4056.17 | 46.77 |

**Table A5.**
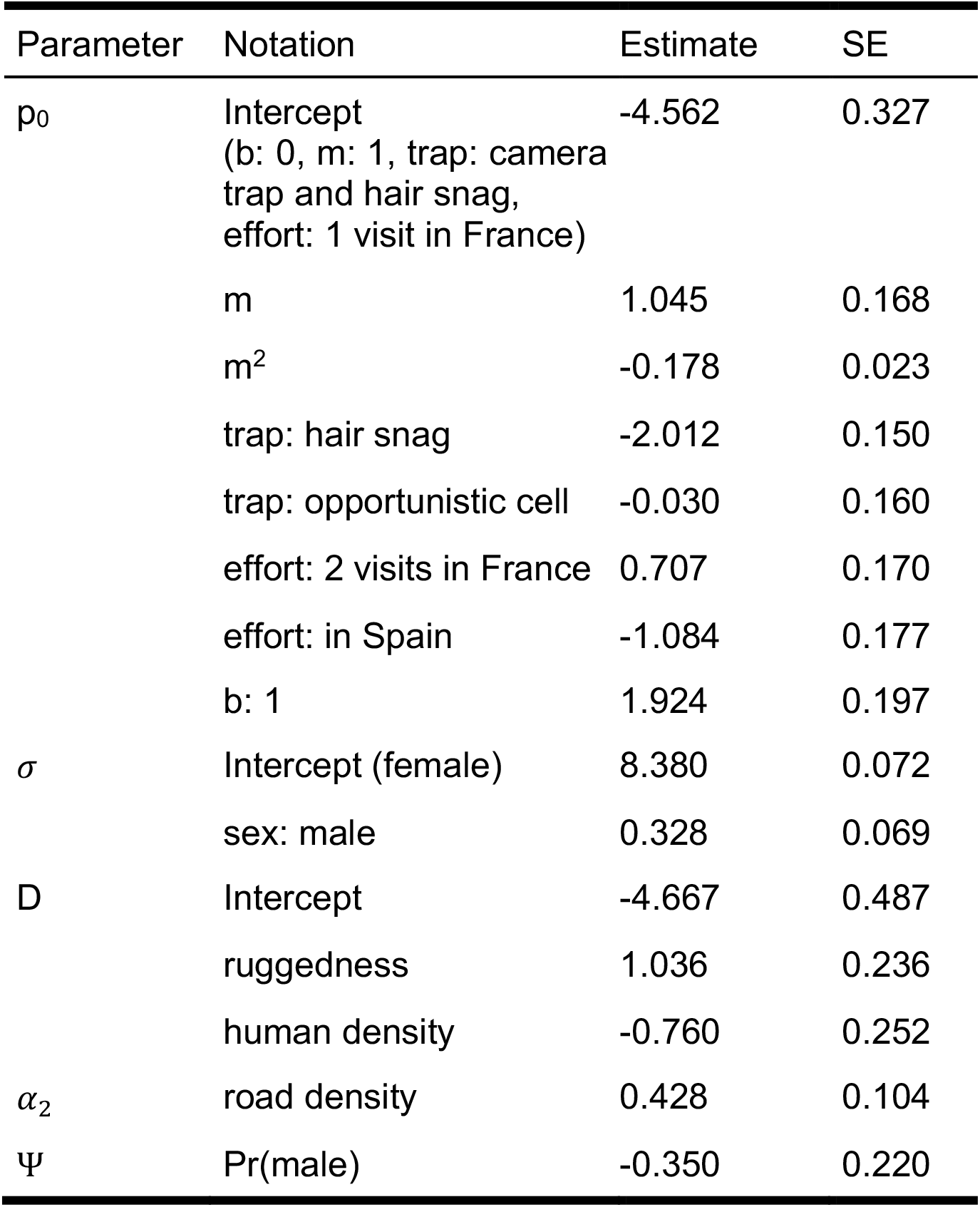
Maximum likelihood estimates (MLE) and standard errors at the end of the hierarchical model selection. In the top model, D is the density of bears and considers an additive effect of the ruggedness and the human density. The baseline detection probability p_0_ is a linear function of the behavioral response to traps (b), the quadratic effect of the month (m + m^2^), the type of trap (trap) (i.e. hair snag alone, hair snag combined with a camera trap, or the center of the a active cell in the opportunistic grid at 5 km resolution), and the search effort (effort) (i.e. depends of the country where the trap is located and the number of visits per month). The scale parameter σ is sex dependent. The resistance parameter α_2_ depends of the road density (road density) per cell at 2.5km resolution. ((D ~ ruggedness + human density) (p_0_ ~ b + m + m^2^ + trap + effort) (σ ~ sex) (α_2_~ 1 − road density))

| Parameter | Notation | Estimate | SE |
| --- | --- | --- | --- |
| $p_0$ | Intercept<br>(b: 0, m: 1, trap: camera trap and hair snag, effort: 1 visit in France) | -4.562 | 0.327 |
|  | m | 1.045 | 0.168 |
| | $m^2$ | -0.178 | 0.023 |
|  | trap: hair snag | -2.012 | 0.150 |
|  | trap: opportunistic cell | -0.030 | 0.160 |
|  | effort: 2 visits in France | 0.707 | 0.170 |
|  | effort: in Spain | -1.084 | 0.177 |
|  | b: 1 | 1.924 | 0.197 |
| $\sigma$ | Intercept (female) | 8.380 | 0.072 |
|  | sex: male | 0.328 | 0.069 |
| $D$ | Intercept | -4.667 | 0.487 |
|  | ruggedness | 1.036 | 0.236 |
|  | human density | -0.760 | 0.252 |
| $\alpha_2$ | road density | 0.428 | 0.104 |
| $\Psi$ | Pr(male) | -0.350 | 0.220 |

**Table A6.** Comparison of 4 representations of the road network on the resistance parameter (alpha2) to the Euclidean model, ranking according to their AIC and considering only data from French monitoring. K is the number of parameters, AIC is the Akaike Information Criterion and ΔAIC is the difference between the AIC of each model and the model with the lowest AIC. The model considered is ((D ~ ruggedness + human density) (p_0_ ~ b + m + m^2^ + trap + effort + session) (σ ~ sex) (α_2_ ~ 1 − road))

| road | K | AIC | $\Delta AIC$ |
| --- | --- | --- | --- |
| Density of all roads | 16 | 3243 | 0 |
| Density of motorway and primary roads | 16 | 3246 | 3 |
| Distance to motorway and primary roads | 16 | 3271 | 28 |
| 1 (Euclidean) | 15 | 3329 | 86 |

**Figure A1.**
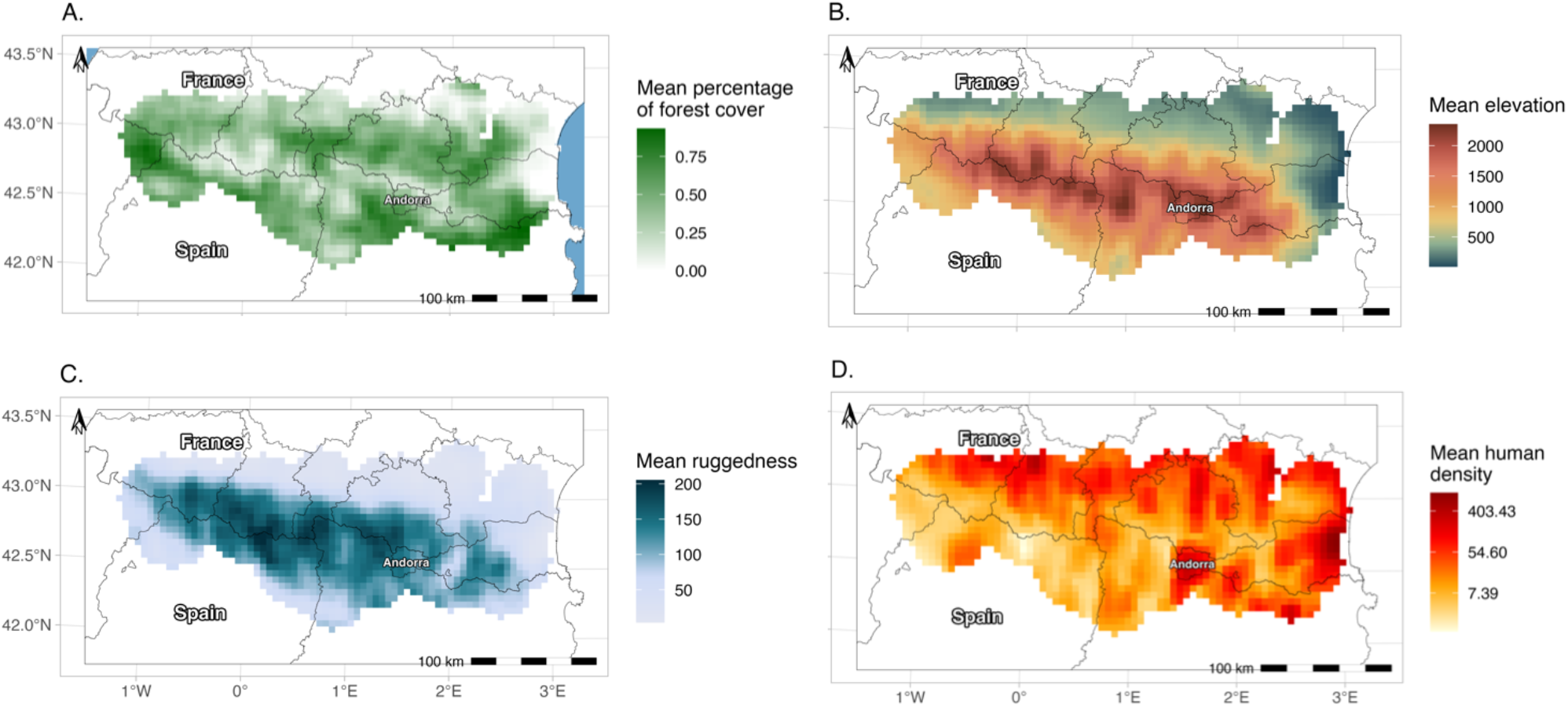
Maps of landscape variables in the Pyrenees (France, Spain, Andorra) at 5 km resolution on the study area. (A) Forest cover (percentage of forest cover averaged on 200 km^2^). (B) Elevation (elevation averaged on 200 km^2^). (C) Ruggedness (ruggedness averaged on 200 km^2^). (D) Human density (human density averaged over 200 km^2^ on a logarithmic scale)

**Figure A2.**
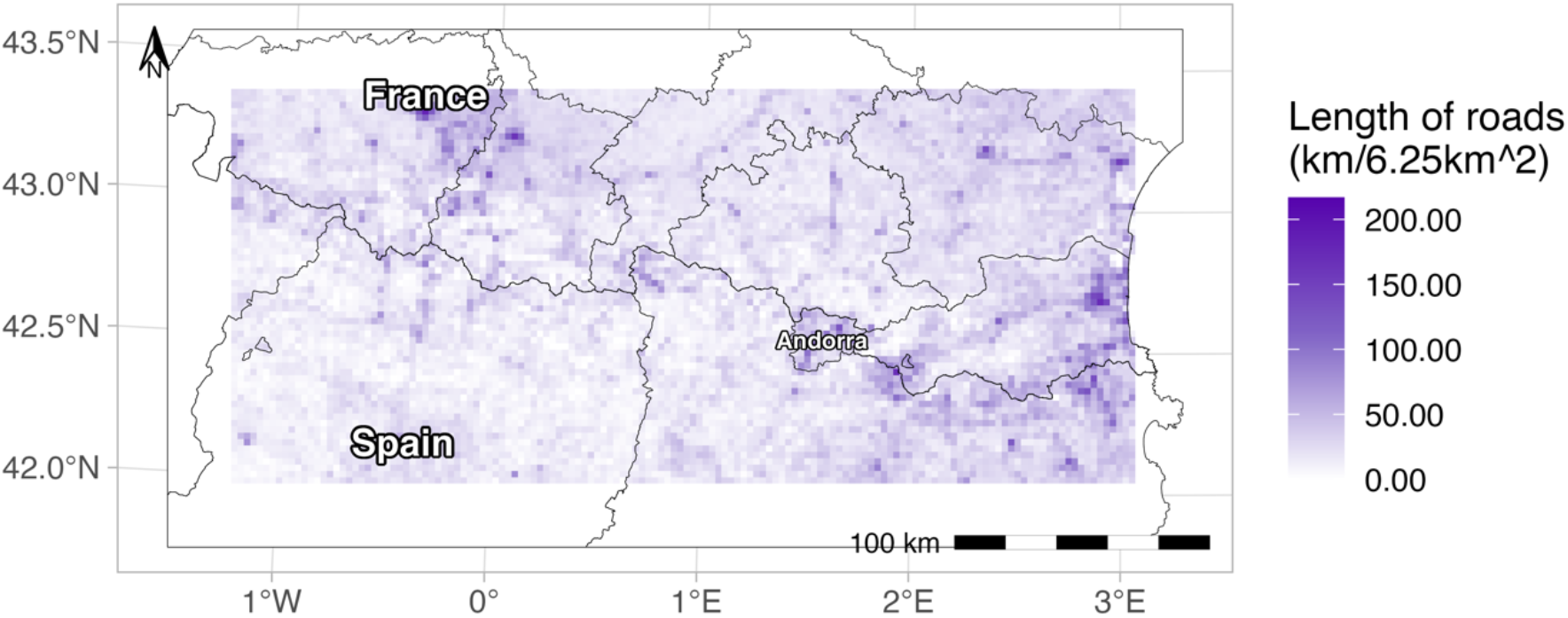
Map of road density in the Pyrenees (France, Spain, Andorra) at 2.5 km resolution on the study area. All roads define in OpenStreetMap are considered.

**Figure A3:**
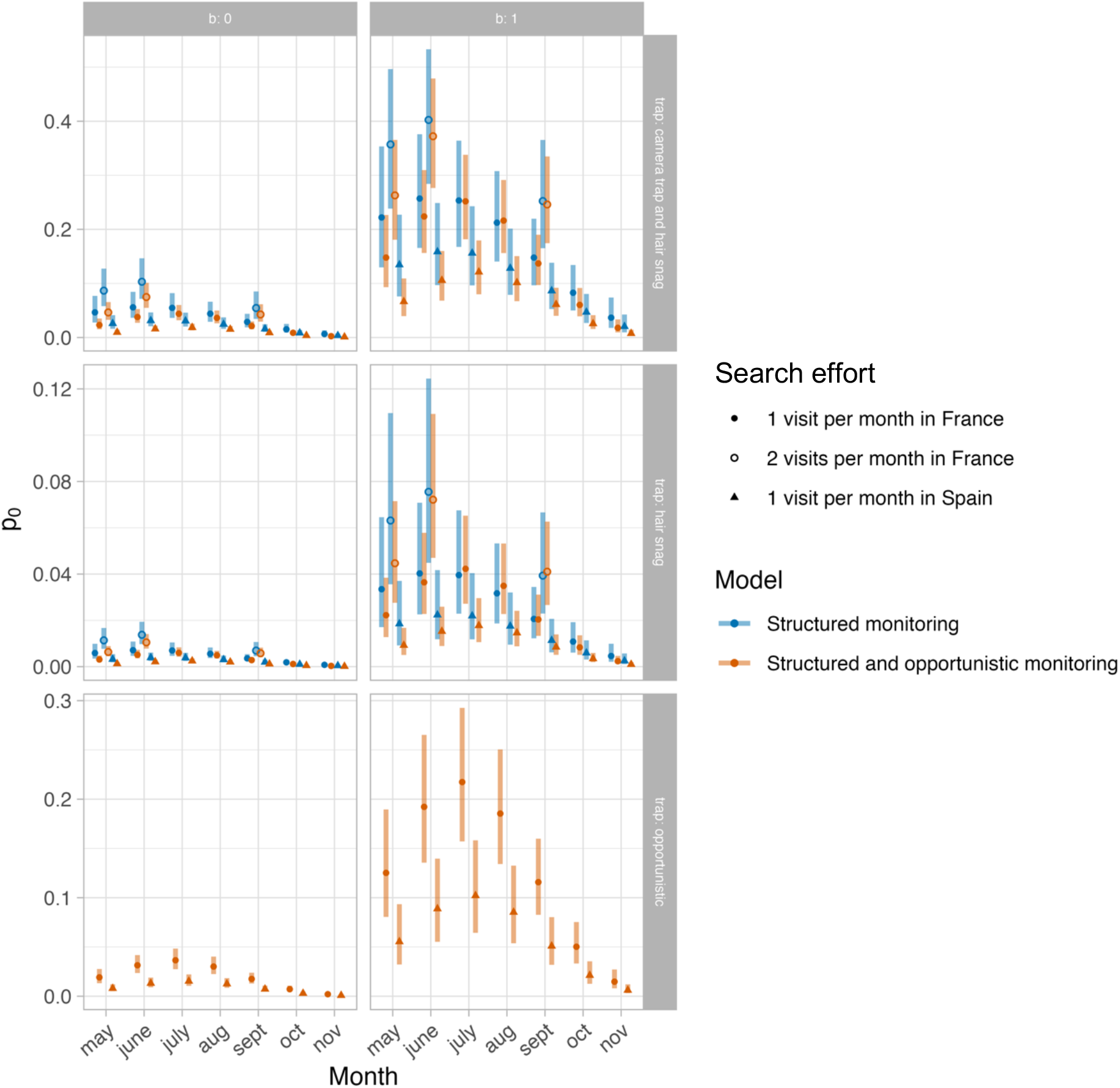
Baseline detection probability (p_0_) in function of the month of the year (from May to November), according to the type of trap (hair snag alone for centered panels or hair snag combined with a camera trap for top panels and opportunistic traps for bottom panels) and if the individual has never been detected (b=0 ; left panels) or was at least detected once (b=1; right panels). Two models are considered: A model with only data from structured monitoring (in blue) and a model with both structured and opportunistic monitoring (in orange). The points represent the estimated values and the bars the 95% confidence interval, when the trap was visited only once per month in France (filled point), twice per month in France (empty point) or once per month in Spain (triangle).

**Figure A4.**
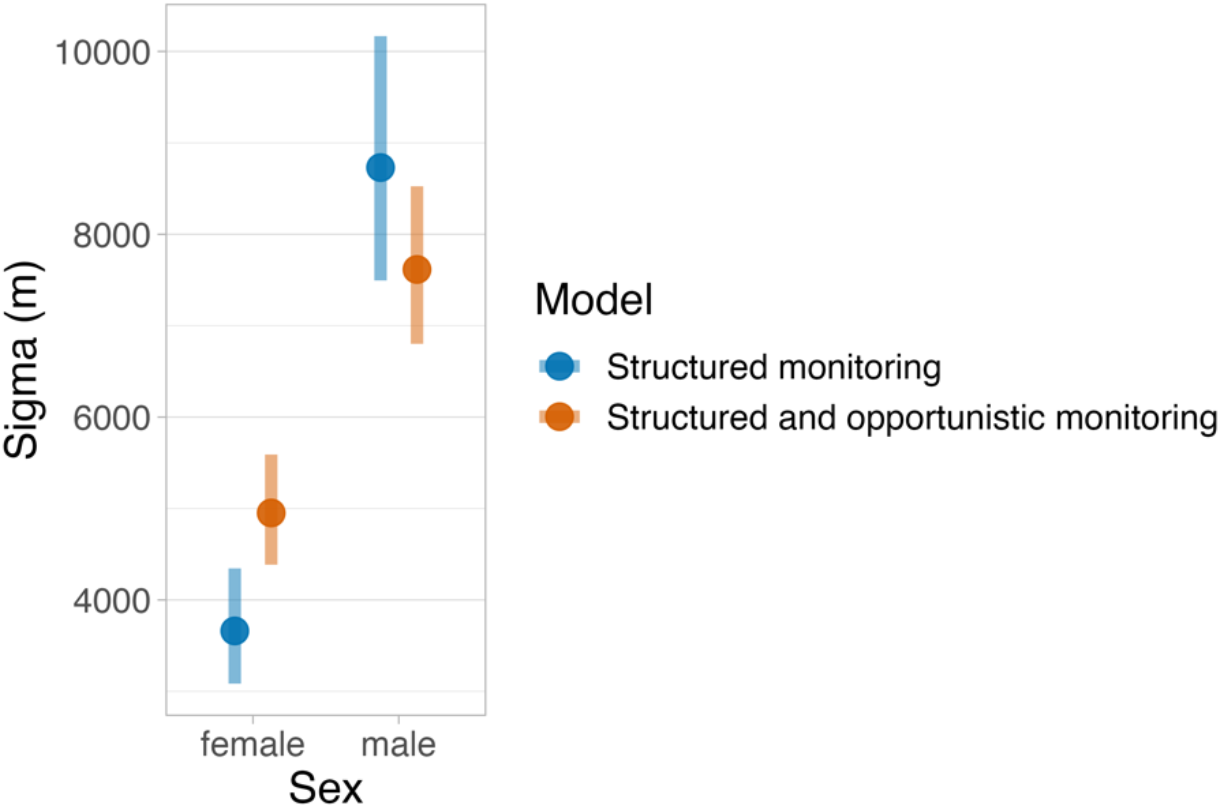
Scale parameter (σ) in meters for females and males. The estimates of two models are compared ((D ~ 1) (p_0_ ~ b + m + m^2^ + trap + effort + session) (σ ~ sex))). A model with only data from structured monitoring (in blue) and a model with both structured and opportunistic monitoring (in orange).

**Figure A5.**
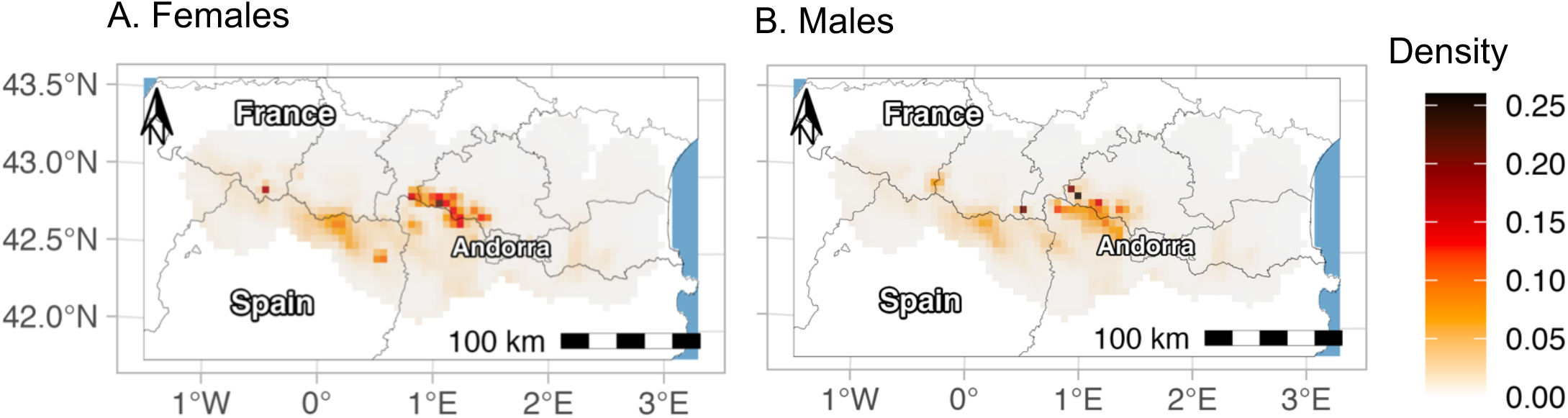
Maps of the realized density of female (A) and male (B) Pyrenean brown bears (in number of bears per km^2^) in 2019. The grey shaded zone represents the study area.

## Home range size

Here, we mapped variation of brown bear home range size in the Pyrenees for females and males and we quantified it in function of road density. For each cell su of the study area S, we separately computed the home range size from the 95% kernel of the utilization distribution for females and males. We supposed that an activity center was present in each cell si in S. To avoid edge effect, that is home ranges are smaller near the edge, we deleted a buffer of 25 km around the study area. Then, we modeled this estimated home range size in km^2^ in function of road density (in km per cell of 6.25 km^2^) in the cell of the activity center. Because home ranges sizes were positives, we used a Poisson generalized linear model. The 95% confidence intervals were estimated with a parametric method with the package ciTools (version 0.6.1) from the expression: 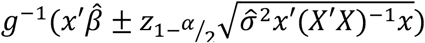

**Figure A6.**
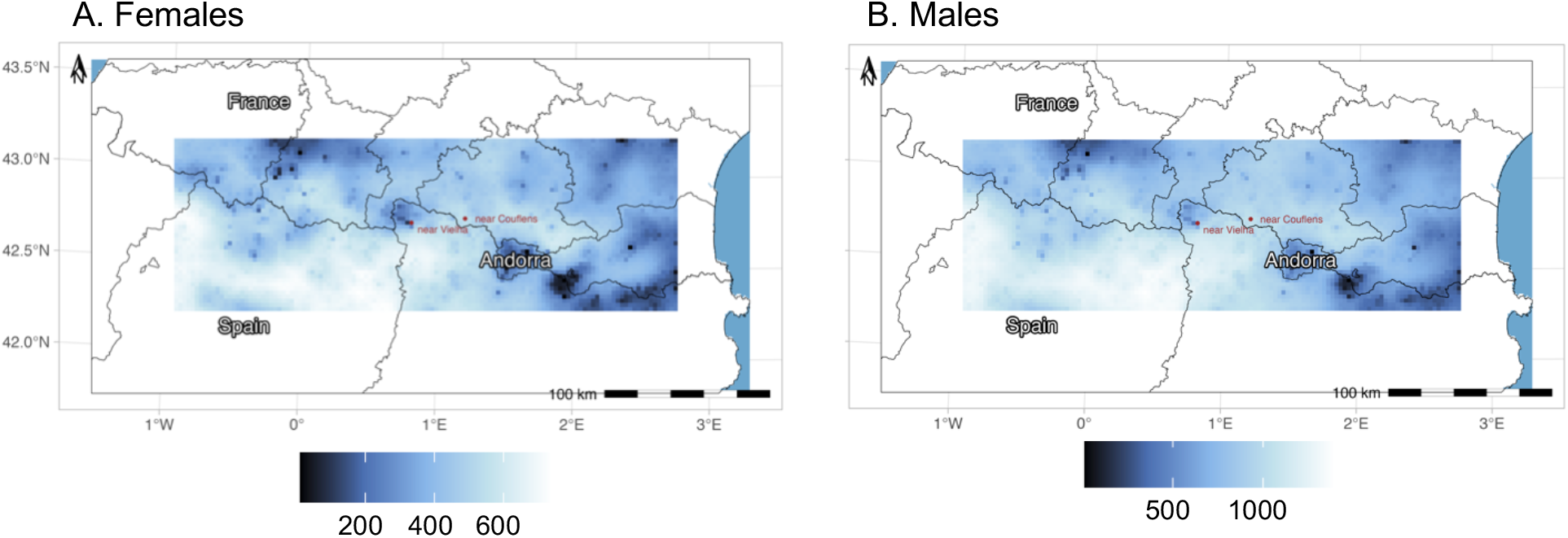
Maps of the home range size of female (A) and male (B) Pyrenean brown bears (in km^2^) computed from the 95% kernel of the utilization distribution at a resolution of 6.25 km^2^. The red point near Couflens is representative of a zone with a low road density (1.38 km/km^2^) and the red point near Vielha is representative of a zone with a high density (8.29 km/km^2^).

**Figure A7.**
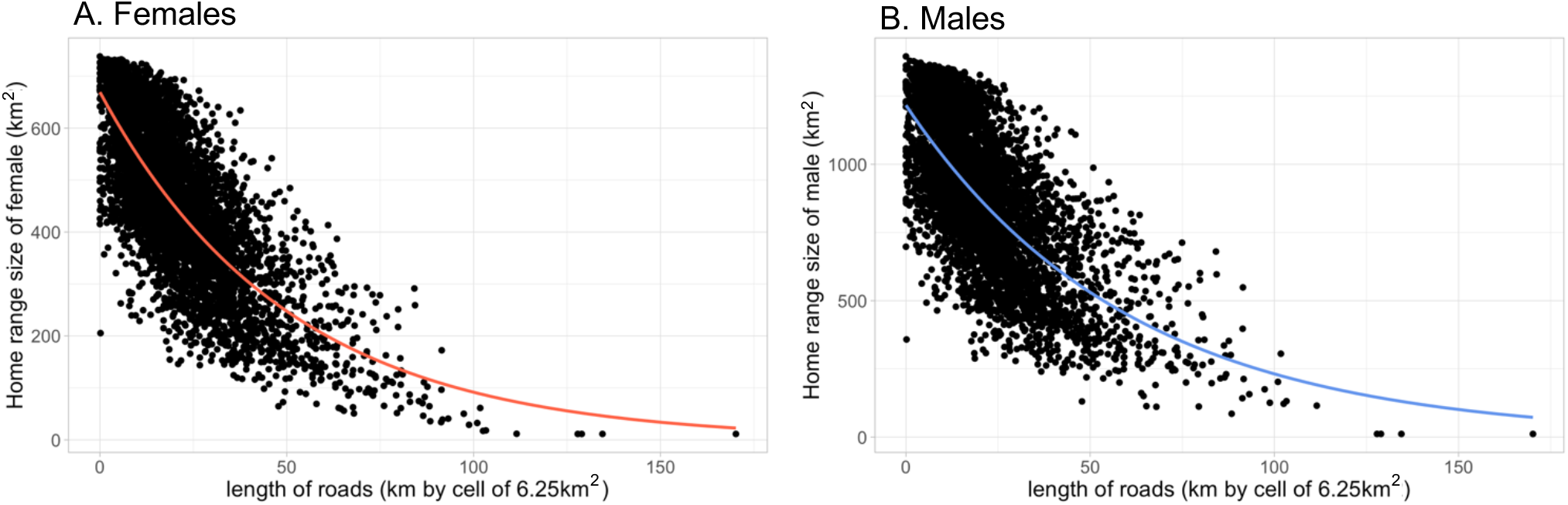
Home range size of female (A) and male (B) Pyrenean brown bears (in km^2^) estimated from the 95% kernel of the utilization distribution in function of road density at the activity center (in km per cell of 6.25 km^2^). The curves represent the estimated values with a Poisson generalized linear model and the shaded zone the confidence interval at 95%.

